# Three-dimensional visualization of blood vessels in human gliomas based on tissue clearing and deep learning

**DOI:** 10.1101/2023.10.31.564955

**Authors:** Xinyue Wang, Xiaodu Yang, Dian He, Yunhao Luo, Qiyuan Huang, Ting Li, Zhaoyu Ye, Chun Ye, Minglin Zhang, Hei Ming Lai, Yingying Xu, Haitao Sun

## Abstract

Gliomas, with their intricate and aggressive nature, call for a detailed visualization of their vasculature. While many studies lean towards 2D imaging of thin sections, this method often overlooks the full spatial heterogeneity inherent to tumors. To overcome this limitation, our study melded state-of-the-art techniques, encompassing tissue clearing technology, 3D confocal microscopy imaging, and deep learning-aided vessel extraction, resulting in a comprehensive 3D visualization of glioma vasculature in intact human tissue. Specifically, we treated formalin-fixed thick human glioma tissue sections (500 μ m) with OPTIClear for transparency and subsequently performed immunofluorescent labeling using CD31. Using confocal microscopy, we obtained 3D images of the glioma vasculature. For vessel extraction, we employed a specialized 3D U-Net, enriched with image preprocessing and post-processing methods, and benchmarked its performance against the Imaris software. Our findings indicated that OPTIClear-enabled tissue clearing yielded a holistic 3D representation of immunolabeled vessels in clinical human glioma samples. Impressively, our deep learning technique outshined the traditional Imaris approach in terms of accuracy and efficiency in vessel extraction. Further, discernible variations in vascular metrics, such as mean diameter, branching point count, and volume ratio, were observed between low-grade and high-grade gliomas. In essence, our innovative blend of tissue clearing and deep learning not only enables enhanced 3D visualization of human glioma vasculature but also underscores morphological disparities across glioma grades, potentially influencing pathological grading, therapeutic strategies, and prognostic evaluations.

## 1 Introduction

Glioma represents the predominant primary brain tumor, constituting about 70-80% of all malignant intracranial tumors[33, 35, 36]. Glioblastoma, classified as WHO grade IV, emerges as the most aggressive glioma subtype[13, 26]. Alarmingly, less than 3% of patients endure beyond 12-15 months due to its rapid malignancy[34]. Its intricate inter- and intra-tumoral heterogeneity coupled with its convoluted microenvironment interactions amplify the challenge in deciphering underlying mechanisms[3, 4, 20, 32].

Angiogenesis plays a key role in tumor growth by providing oxygen and nutrients for tumor cells[54]. Glioblastomas manifest with pronounced vasculature but compromised blood-brain barrier (BBB) structural integrity[18].This vascular profusion results in erratic permselectivity, perfusion, elevated interstitial fluid pressure, rampant hypoxia, necrosis, and edema [18]. Hypoxic conditions induce gene expression shifts in tumor cells, bolstering cell survivability and apoptosis resistance [54]. And high interstitial fluid pressure can prevent drug penetration into tumor tissue[15]. In summary, the morphological structure of glioma vessels is closely related to their function, so further studies of their morphological features are crucial to provide further insight into their function. Amid the myriad regulatory mechanisms steering glioma angiogenesis[41, 46, 53], these vessels exhibit pronounced phenotypic heterogeneity, especially concerning their morphology, wall integrity, and angiogenic patterns[1, 10, 47]. Such variations influence glioma progression and prognosis. Historical vascular research largely hinged on optical microscopy of micron-thick sections, yielding two-dimensional (2D) imagery. This 2D approach significantly restricts imaging depth. Breakthroughs in tissue clearing now permit deep tissue labeling and whole-tissue specimen processing without sectioning, ushering in a new era of three-dimensional (3D) tissue imaging via advanced microscopy[27, 57].Numerous tissue clearing methodologies, such as CLARITY and iDISCO/uDISCO, are now available tailored to specific cells or tissues[6, 27, 37, 42, 57]. Our team has previously pioneered the OPTIClear solution for the human brain, designed to exclusively modulate tissue optical properties without compromising structural or molecular integrity.This reveals neural structures and the 3D vascular architecture when coupled with 3D reconstruction technology[22]. While animal studies have correlated 3D vascular spatial heterogeneity with glioma cell invasiveness[21], human glioma tissue studies remain scant.

In order to present fine vascular structure and capture the morphological changes, the vessels need to be extracted from the complex background and quantitatively characterized. Currently, vessel extraction and analysis in cleared tissue images mostly rely on simple filter-based image processing algorithms [19, 31, 56] or interactive software, for example, Imaris[21, 24, 28]. It is difficult for them to realize customized and automatic extraction and computation of high-throughput vascular data. Moreover, the accuracy of vessel segmentation is far from satisfactory and often relies on subsequent manual adjustment. Nowadays, deep learning algorithms, which are the mainstream in artificial intelligence, provide a new solution for the vessel segmentation of 3D cleared tissue images[23, 49]. They can achieve fast, automatic, and accurate extraction of the cleared tissue data, which is of great significance for achieving high-throughput and bias-free object extraction and analysis. 3D U-Net is one of the most classic neural networks for segmentation of 3D biomedical images [7], and is able to cope with many challenging issues, such as low signal-to-noise ratios, sparsity in annotation, and insufficient amount of training data[11].

In this research, we amalgamate tissue clearing with deep learning to elucidate the 3D architecture and heterogeneity of glioma vessels in human samples. Using the OPTIClear method on 500 μm thick specimens, we achieved 3D imaging of glioma vessels. Subsequently, a customized 3D U-Net combined with image preprocessing and post-processing methods was built to automatically extract vessels from variable and complex background. Finally, we extracted vascular features and conducted qualitative and quantitative analysis, which demonstrated a distinct heterogeneity in the vasculature of gliomas across various grades..

## 2 Materials and Methods

### 2.1 Preparation of human Samples

The human glioma samples used in this study were obtained during neurosurgeries at the Zhujiang Hospital. Based on the WHO Classification of gliomas, we selected tumors clinically diagnosed as low-grade (grade II astroglioma) and high-grade gliomas (grade IV glioblastoma). Both tumors were surgically resected from the right frontal region of the brain. Furthermore, normal brain tissues from the same region in non-tumorous human brains were used as controls. All the samples were cleaned with normal saline before fixed in 10% neutral buffered formalin (NBF; Wexis, Guangzhou, China) for at least 5-7 days at 4 ℃.

### 2.2 OPTIClear Optical Clearing and Immunolabeling

Tissue blocks of appropriate size were cut from the fixed tissue, and the excess fixative was rinsed with phosphate buffered saline (PBS; Solarbio, Beijing, China). Then the tissue block placed in homemade aluminum molds was flooded with 4% (vol/vol) agarose (Sigma, American) and solidified on ice. Next, the tissue block embedded in agarose gel was cut into 500 μm thick sections with a vibratome (Koster, American).

After removal of excess fixative with water and removal of excess agarose with tweezers, processed tissue blocks were immersed in Sodium borate clearing (SBC; 4% (wt/vol) SDS dissolved in 0.2M sodium borate, pH 8.5) for delipidation for 3-5 days at 37-55℃. Next, the tissue was washed in phosphate-buffered saline (PBS) with Triton X-100 (PBST; 0.3% Triton X-100 (vol/vol) and 0.01% sodium azide (wt/vol)) at 37 ℃ (3×2h). The tissue was then immunostained with primary antibody (rabbit anti-CD31 antibody; Abcam, Shanghai, China) in PBST at 37 ℃ for at least 2 days. Primary antibodies were removed by washing with PBST (3×2h) and then left overnight. Sections were incubated with secondary antibodies (Goat anti-rabbit antibody, Alexa Fluor^®^ 633 conjugate; sigma, American) overnight, protected from light. After washing off secondary antibodies with PBST(3×2h, at 37℃) and left overnight, sections were incubated with OPTIClear solution (an aqueous solution consisting of 20% (wt/vol) N-methylglucamine, 32% (wt/vol) iohexol and 20.48% (vol/vol) 2,2’-thiodiethanol (TDE), with a pH between 7 to 8 adjusted with hydrochloric acid) for 6 h to clear the tissue.

### 2.3 Immunohistochemistry and immunofluorescence staining of FFPE sections

Tissue blocks of appropriate size were cut from the fixed tissue and cut into 4 μm-thick slices after embedded in paraffin. Next, deparaffinization, rehydration, blocking of endogenous peroxidase (30 min at RT in 3% H_2_O_2_), antigen retrieval (two boiling/standing cycles for 20 min in sodium citrate solution, pH 6.0), and blocking with 5% normal goat serum (Histova, Beijing, China) for 30 min at 37℃ were performed according to the standard immunohistochemistry staining protocol. Primary antibodies (rabbit anti-CD31 antibody; Abcam, Shanghai, China) were applied on slices overnight at 4℃ and secondary antibodies (Goat Anti-Rabbit IgG Antibody, (H+L) HRP conjugate; sigma, American) were added to incubate for 1h at 37℃. Then freshly prepared DAB chromogenic solution (Servicebio #G1212-200T) was added and the color reaction was observed under the microscope (Nikon E100, Japan), terminated in time with distilled water. Nuclei were counterstained with hematoxylin (Solarbio #G1004) for 3-5min and then the sections were dipped in 1% hydrochloric acid in alcohol for 3s for differentiation. PBS was added dropwise on the tissue section to return to blue for 1min. Then gradient alcohol dehydration, xylene hyalinization and neutral gum sealing were performed. A bright-field panoramic digital slide scanner (3D HISTECH, Hungary) was used to view the slides.

When immunofluorescence staining performed, the procedures described above were repeated until the primary antibody incubation step. Then secondary antibodies (Goat anti-rabbit antibody, Alexa Fluor^®^ 633 conjugate; sigma, American) were added to incubate for 1h at 37℃. DAPI - containing mounting media (Fluoroshield Mounting Medium With DAPI, Abcam #ab104139) were used for mounting with coverslip and samples were analyzed using Leica SP8 confocal laser scanning microscope.

### 2.4 Confocal Microscope Imaging

All cleared tissues were mounted on 60-mm cell and tissue culture dishes wetted with OPTIClear solution (about 200-400 μL) under the microscopy. Images were obtained and scanned in 3D along the Z-axis with a Leica SP8 confocal microscope. Thus, a volume data, which refers to the 3D image of one region of interest in a sample, was collected. To achieve good contrast with minimal detection of autofluorescence signal, a detection spectral range of 570-700 nm was chosen.

### 2.5 3D visualization and modeling of vasculature in Imaris

Immunohistochemical staining of FFPE sections was visualized using LAS X (Leica software). Original 3D immunofluorescence images and 3D segmentation masks acquired by deep learning method, were visualized using Imaris 9.0.1 (Bitplane software). The former was visualized with maximum intensity projection (MIP) mode and the latter was visualized with blend mode to show the 3D effect. In the overlapping images of them, we replaced the blend with MIP mode and adjusted its transparency.

The Filament Tracer tool in Imaris, designed for filament-like structures, was used to construct the vascular computational models for vessels extraction with Imaris. In order to reduce the burden of computational modeling and obtain better models, the images with low signal-to-noise ratios were preprocessed with ‘Background Subtraction’ option of Imaris. Then, the filament models were constructed with a threshold-based algorithm and presented in Cone style with green color. The transparency of models was adjusted to better show the overlap of the immunofluorescence volumes and filament models.

### 2.6 Vessel extraction with deep learning network

After obtaining 3D images, we used a deep learning network to automatically extract vessels from complex tissue background. The raw images were preprocessed before fed into the deep learning network, and the segmentation results of the network were post-processed to obtain more accurate results.

#### 2.6.1 Dataset and preprocessing

Due to the difficulty of annotation, fourteen representative subvolumes from the whole volumes were selected for network training and model selection, and the ground truth was manually annotated in one slice for each ten slices using ITK-snap. To evaluate the segmentation performance objectively, we used two-fold cross validation. The subvolumes were divided into two sets, one composed of seven subvolumes from four patients and the other composed of the remaining seven from the other four patients. Both of the two sets have various vascular appearances and tumor grades.

We preprocessed the 3D images with contrast limited adaptive histogram equalization algorithm[39] and rolling ball background subtraction algorithm[48] to enhance weak vessels and suppress strong background. Then, the data was normalized to unify the voxel intensity.

#### 2.6.2 Deep learning network architecture, training, and inference

Motivated by U-Net’s great success in segmenting various biomedical images[7, 9, 17, 38], we trained an improved 3D U-Net (Figure 1) for glioma vessel segmentation. It consists of a contracting path to capture context information and a symmetric expansive path to recover precise location information, called encoder and decoder, respectively. The information from the encoder is combined into the decoder via skip connections. The architecture has 6 resolution stages and each stage consists of two-layer convolution blocks. Different from the original 3D U-Net, in the convolution block, we replaced the batch normalization with instance normalization[51] due to the small batch size of 2. Furthermore, we replaced ReLU with leaky ReLU[29] as the activation function. With the stride and kernel size which were adapted to our image size and voxel size, downsampling was conducted with strided convolutions in the encoder and upsampling was conducted with convolutions transposed in the decoder. The input of network was a preprocessed 3D image sampled with a size of 48×192×192 and the output was a matrix of probability with same size of the input image. The matrix was then binarized using a threshold of 0.5 to yield the segmentation mask.

**Fig. 1.**
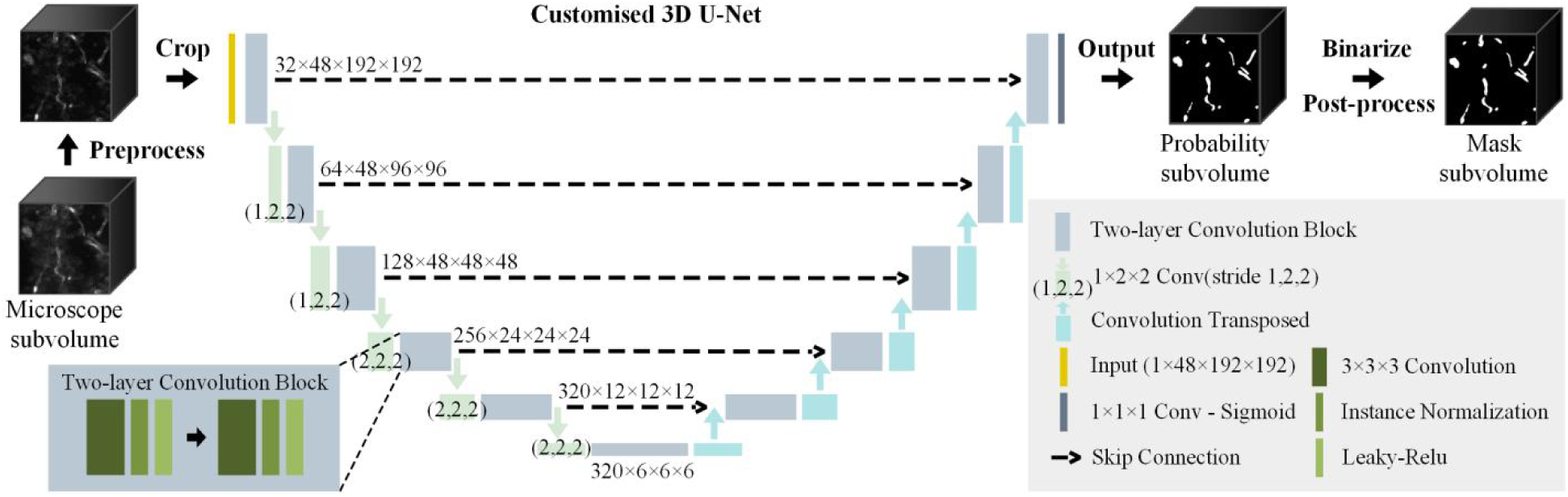
Vessel extraction method based on 3D U-Net. The sizes of feature maps are indicated after the blocks.

The background has much more pixels than the vessels, so dice loss was used as the training loss. The input images were sampled randomly and contain at least one foreground pixel. To increase the diversity of the data and improve the model robustness, we used diverse dynamic data augmentation strategies including random rotation, flipping, gamma transformation, and introduction of Gaussian noise. The training of network was driven by the Adam optimizer with an initial learning rate of 1×10^-4^ and an exponential decay rate of 0.9 for the first moment and 0.99 for the second moment. The training runs for 600 epochs and each epoch has 250 training iterations. The network training was on a single NVIDIA Quadro P6000 GPU using Tensorflow framework and KERAS.

The test images were predicted with a sliding window with the input size. To improve the performance of prediction, we applied data augmentation by flipping along three axes and then got the final prediction by averaging them. Dice coefficient, commonly used in segmentation tasks to measure the similarity of two images, was used as the evaluation metric for the vessel extraction. It can be calculated by the following equation:

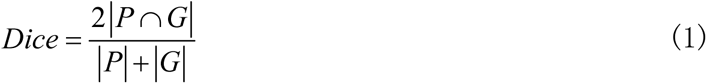

Where *P* presents the pixel set in the prediction images and the *G* presents the pixel set in the ground truth.

#### 2.6.3 Post-processing

To further improve the segmentation results, we performed 3D connected component analysis and morphological operations on the segmentation masks. Firstly, the images were denoised by removing the components which is smaller than a certain volume value *v*. Then, the denoised images were processed by 3D closing operation in mathematical morphology to connect vessel segments and fill small holes. The closing operation consists in the succession of a dilation and an erosion operation, which is repeated with *n* iteration times. Finally, we applied 3D filling holes function in mathematical morphology to the closed images. The parameters *v* and *n* were determined by grid search based on the segmentation performance.

### 2.7 Feature quantification and statistical analysis

To investigate the spatial heterogeneity of blood vessels in different grades of human glioma, we applied the above segmentation method in the volumes of all the samples, and extracted the vascular morphological features based on the segmentation results[49]. The vascular features included mean diameter along the full length (um), number of branching points per unit volume (count/mm^3^), volume ratio, surface area ratio, and total vessel length per unit volume (mm/mm^3^). Volume ratio is the ratio of the total voxel number of vascular volume to the total voxel number of the whole volume. Surface area ratio is the ratio of the total voxel number of vascular surface to the total voxel number of the whole volume. They were firstly calculated in the voxel space and then transformed to the anatomical space by introducing the information of voxel size.

Subsequently, statistical analysis was performed using GraphPad Prism 8.0.1 (GraphPad Software). Data were expressed as mean ± SEM. For the two groups that were not significantly different in three group analysis (normal, low grade, and high grade groups), we additionally analyzed their difference to present a more comprehensive analysis result. The significance of differences among three groups and each two groups were tested by one-way ANOVA and unpaired t test if the data satisfied normal distribution and homogeneity of variance. Otherwise, Kruskal-Wallis test, and Welch’s correction or Mann-Whitney test were used, respectively. The results were considered statistically different if the *p* value was less than 0.05.

## 3 Result

### 3.1 OPTIClear is suitable for the visualization of immunolabeled blood vessels in clinical samples of human gliomas

First, we cut the glioma tissue blocks into tissue blocks approximately 5000μm×5000μm×500μm using a vibratome, and then the tissue blocks were processed by OPTIClear. Before the clearing treatment, the tissue block was completely opaque, therefore the English letters underneath the culture dish could not be seen. However, after treatment with OPTIClear tissue clearing, the tissue block exhibited improved light transmissibility without light distortion. Visually, the tissue appeared transparent, allowing letters beneath the transparent culture dish to be clearly discerned without any tissue deformation (Figure 2 a). Then, after immunolabeling and fluorescent staining of blood vessels with CD31 antibodies, images were acquired along the Z-axis using a fluorescent confocal microscope (Leica confocal microscope SP8). The scanning depth was determined by factors including the microscope lens’s working distance(HC PL APO 20×/0.75 CS2, WD=0.62 mm), the tissue block’s thickness, and the slide’s thickness. Therefore, we selected the deep area of the tissue with a thickness of about 100 μm-120 μm for 3D scanning and imaging. Imaris software was used to perform 3D visualization of the images, and the fluorescence images of the intact transparent tissue were obtained. In the high-resolution 3D image, we can not only visualize the blood vessels from various spatial perspectives, but also clearly identify the 3D structure of the vascular tree, including all levels of vascular bifurcation, vessel alignment and other detailed structure (Figure 2 b). Obvious fenestration of endothelial cells can be observed in glioblastoma, suggesting incomplete vessel walls. The above results prove that clearing of human glioma samples using OPTIClear in combination with immunolabeling of glioma vessels with anti-CD31 antibodies make it possible to display a detailed view of the complex spatial structure of the vascular network in intact tissue.

**Fig. 2.**
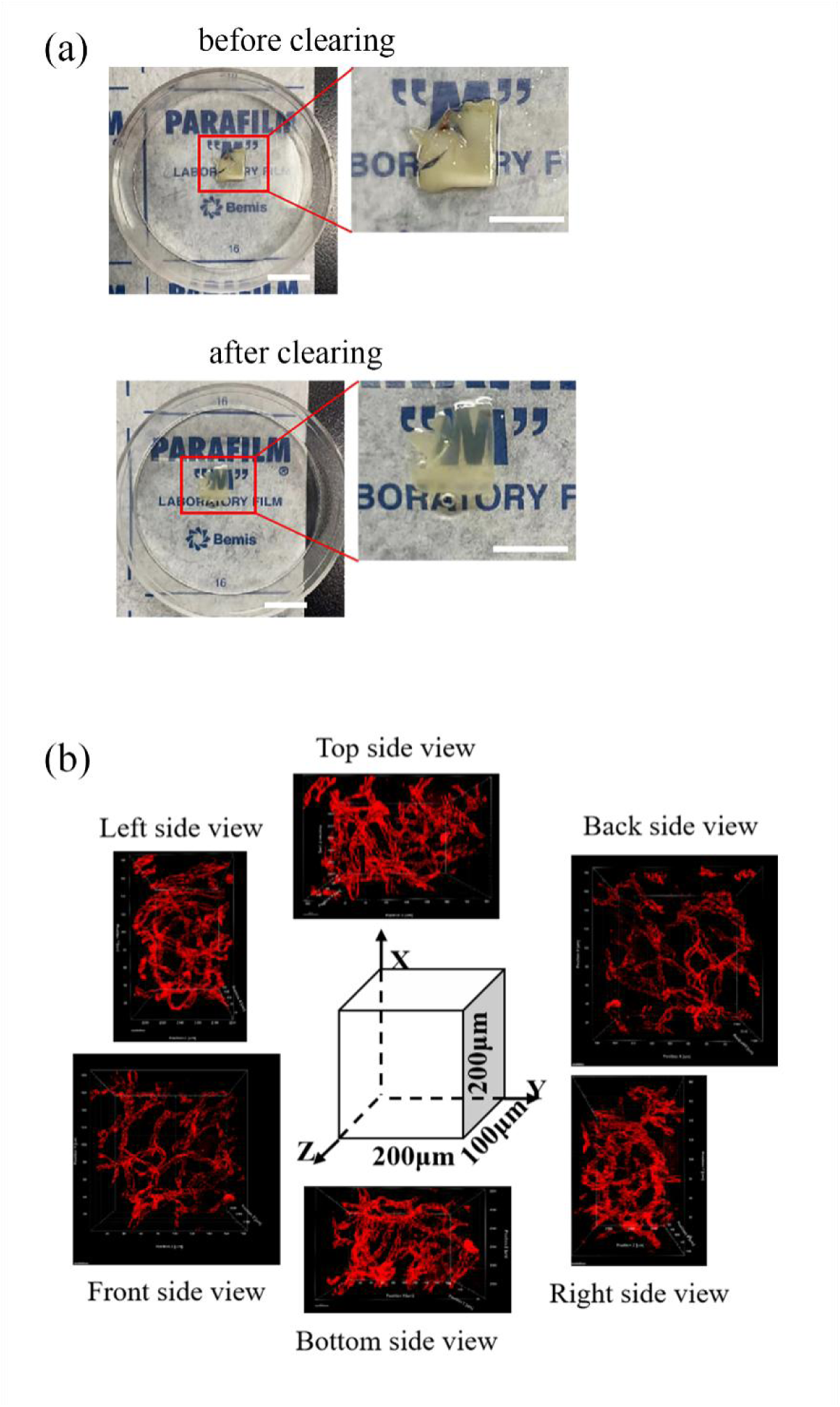
Validation of the applicability of the tissue clearing method for visualization of blood vessels in human gliomas. (a) The clearing result of OPTIClear in human glioma tissue. We can see that before clearing (that is, after fixing with fixative) light cannot penetrate the tissue block, so the text in the background cannot be seen, while after clearing, the light passes through the tissue block and the underlying word can be seen. Scale bar=5mm; (b) 3D visualization of blood vessels in human glioma samples. Anti-CD31 antibody (red) was used to immunolabel the glioma blood vessels, and OPTIClear was used to clear the tissue, the inner 100 μm thick tissue was observed with a confocal microscope and scanned for 3D reconstruction. The 3D reconstructed image of the Z-stack obtained shows a Z-axis thickness of 100 μm. Scale bar=50μm.

### 3.2 3D images better represent the heterogeneity of the glioma vasculature than 2D images

Currently, 2D analysis methods predominate in tumor clinical practice and scientific research, often using immunochemical stained thin tissue paraffin sections for tumor diagnosis and analysis; however, analysis methods reliant on 2D images inherently overlook the intricate 3D spatial anatomy present in intact tissue, potentially bypassing crucial diagnostic or therapeutic insights. In this study, we first observed and analyzed the morphology and distribution of glioma blood vessels in conventional FFPE sections. Results revealed that immunohistochemical staining of these paraffin sections predominantly displayed tubular, spindle, and cluster cross-sectional 2D depictions of blood vessels within a confined region (Figure 3 a, c). This limitation arises from the mechanical sectioning process, where blood vessels are sliced into mere micronic-thin sections. While this conventional 2D pathological approach offers insights into cellular morphology and distribution at certain microscopic scales, it inadequately captures the holistic 3D vascular architecture. Next, we performed tissue clearing and immunolabeling on thick tissue blocks of surgical resection samples from the same patient, and then visualized the 3D images in Imaris software. Obviously, 3D images can present the vascular structure more visually than the 2D images. By utilizing a Z-axis laminar scan of the optical slice at consistent magnification, we transcend the limitations of conventional 2D imaging. The 3D visualizations derived from the cleared tissue offer an intricate and comprehensive perspective on the vascular structure of gliomas from every spatial orientation (Figure 3 b, d).

**Fig. 3.**
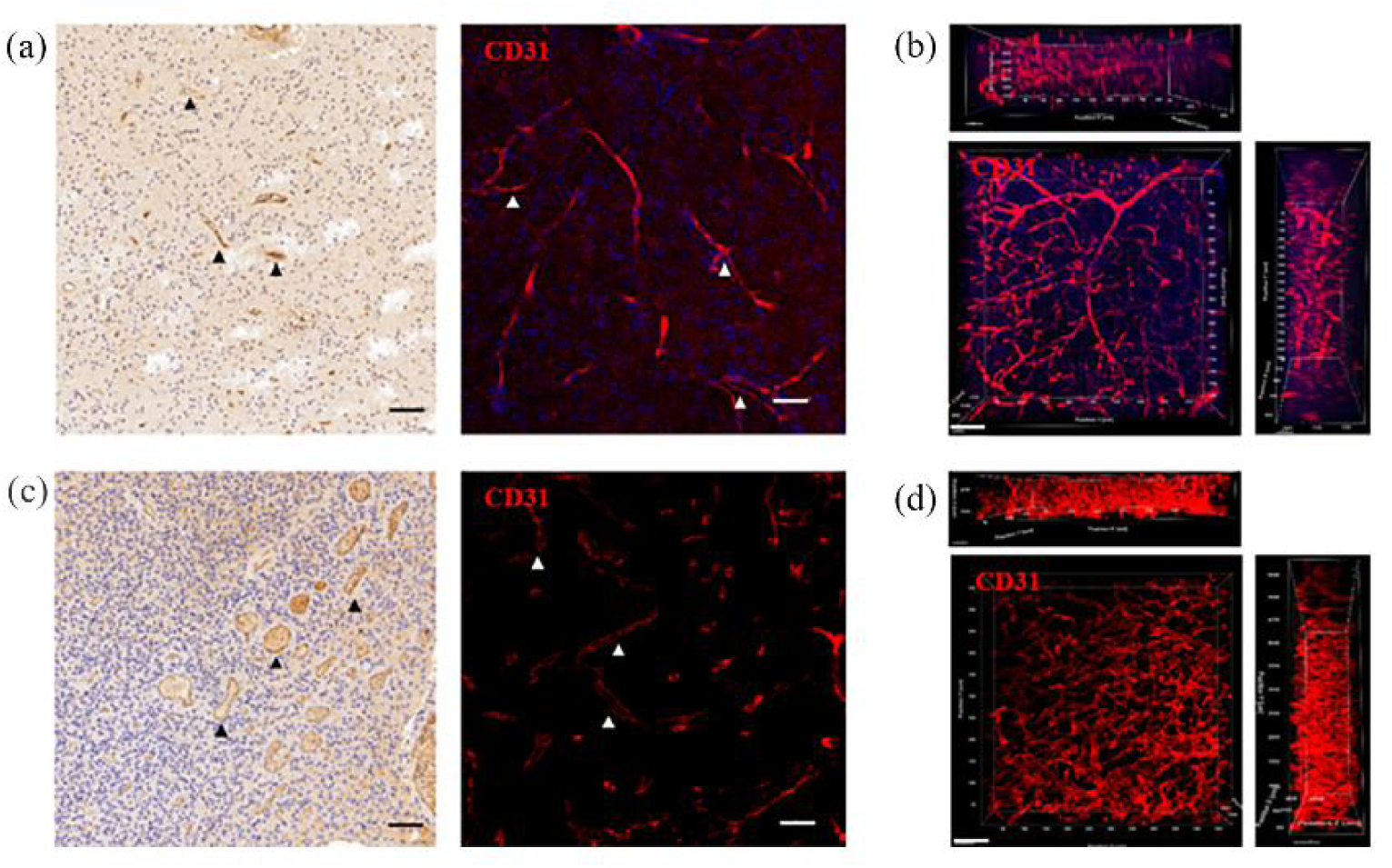
Comparison of 2D and 3D imaging of glioma vessels. (a) Vessels staining of tissue paraffin section of low-grade glioma (grade II), immunohistochemical staining on the left, immunofluorescence staining on the right, the arrow points to the 2D cross-section of the blood vessel, showing a circular or lumen structure; (b) 3D imaging of the clearing of the thick tissue block of low-grade glioma (grade II), marked in red for CD31 vessels staining; (c) Vessels staining of tissue paraffin section of high-grade glioma (grade Ⅳ), immunohistochemical staining on the left, immunofluorescence staining on the right, the arrow points to the 2D cross-section of the blood vessel, showing a circular or lumen structure; (d) 3D imaging of the clearing of the thick tissue block of high-grade glioma (grade Ⅳ), marked in red for CD31 vessels staining. Scale bar=50μm.

Subsequently, based on the 3D images, we delved deeper into visualizing and analyzing the spatial morphological heterogeneity across various glioma grades. Compared with normal cerebrovascular structures, glioma vessels were unevenly distributed and irregularly shaped (Figure 4). The 3D images clearly showed the heterogeneity of vascular spatial morphology, suggesting the significant differences in vascular morphology among normal brain tissue and different grades of gliomas. In addition, we found that glioma blood vessels exhibited disrupted vessel wall. The discontinuous red fluorescent signals presented a broken window-like appearance of the vessel wall and collapsed lumen structures, showing significant difference in the integrity of the vessel wall between low-grade and high-grade gliomas, which is possibly due to the influence of glioma cells. The gray value analysis of CD31 showed consistent levels in low-grade gliomas without large peak-to-trough changes, but sharply changing peaks and valleys in high-grade gliomas, which may reflect more pronounced microscopic breakage of the endothelium in high-grade gliomas than in low-grade gliomas.

**Fig. 4.**
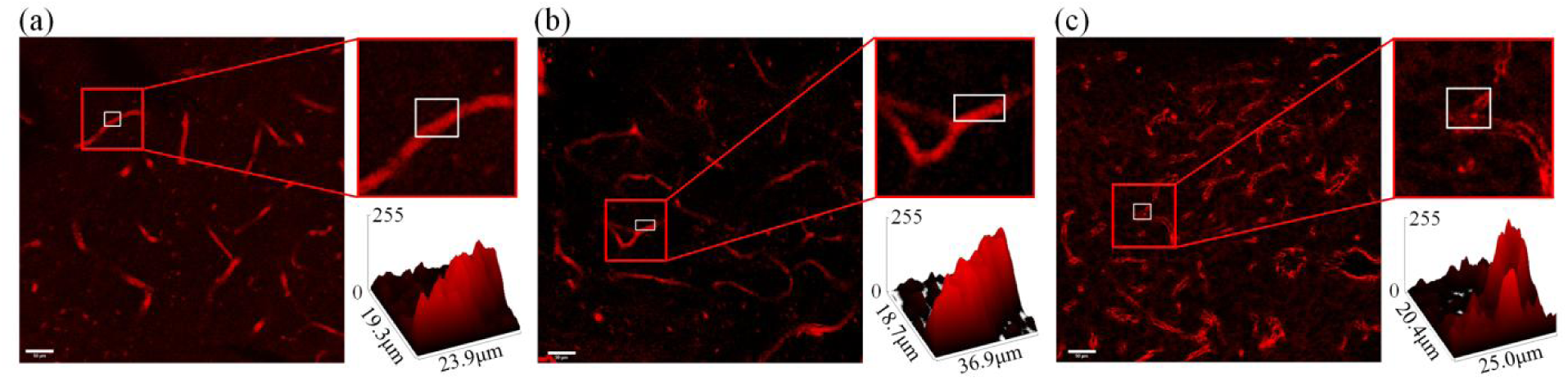
The distribution of CD31 in glioma vessels is uneven and discontinuous, reflecting the rupture of the vessel wall.(a) Cross-sectional view of blood vessels in normal brain tissue (left). Enlarged view (upper right corner) and fluorescence intensity measurement in the blood vessel wall (lower right corner) of the framed area (b) Cross-sectional view of blood vessels in low-grade (grade II) gliomas (left). Enlarged view (upper right corner) and fluorescence intensity measurement in the blood vessel wall (lower right corner) of the framed area ; (c) Cross-sectional view of blood vessels in high-grade (grade IV) gliomas (left). Enlarged view (upper right corner) and fluorescence intensity measurement in the blood vessel wall (lower right corner) of the framed area. Scale bar=50μm.

3.3 Deep learning based algorithms enable more accurate and automated extraction of glioma vessels than the Imaris

In order to better extract the vessel structures and morphological features, we used deep learning-based algorithms to extract glioma vasculature (Figure 5, 6). The average Dice values in 3D U-Net and post-processing are 0.6773 and 0.6867, respectively (Figure 5 a). The representative slices and 3D visualization of subvolumes with different image characteristics, glioma grades, and segmentation qualities were shown in Figure 5 b to give more comprehensive display of the segmentation effects. The first three columns represent 3D raw immunofluorescent subvolumes, its a 2D slice and the pre-processed results with slightly improved image quality, respectively. By comparing our 3D U-Net results and ground truth, we can see that vessels can be extracted by the deep learning method with near manual accuracy. In the post-processed images, some noise was removed and some vessels became more intact as pointed by the red arrows. For the parameter *v* in the denoising operation, we selected the volume value of 290, which is large enough to remove small noises while the Dice value is higher. For the parameter *n* in the closing operation, we selected the value of iteration times as 10, which makes the Dice best. The last column shows the 3D visualization of final segmentation results.

**Fig. 5.**
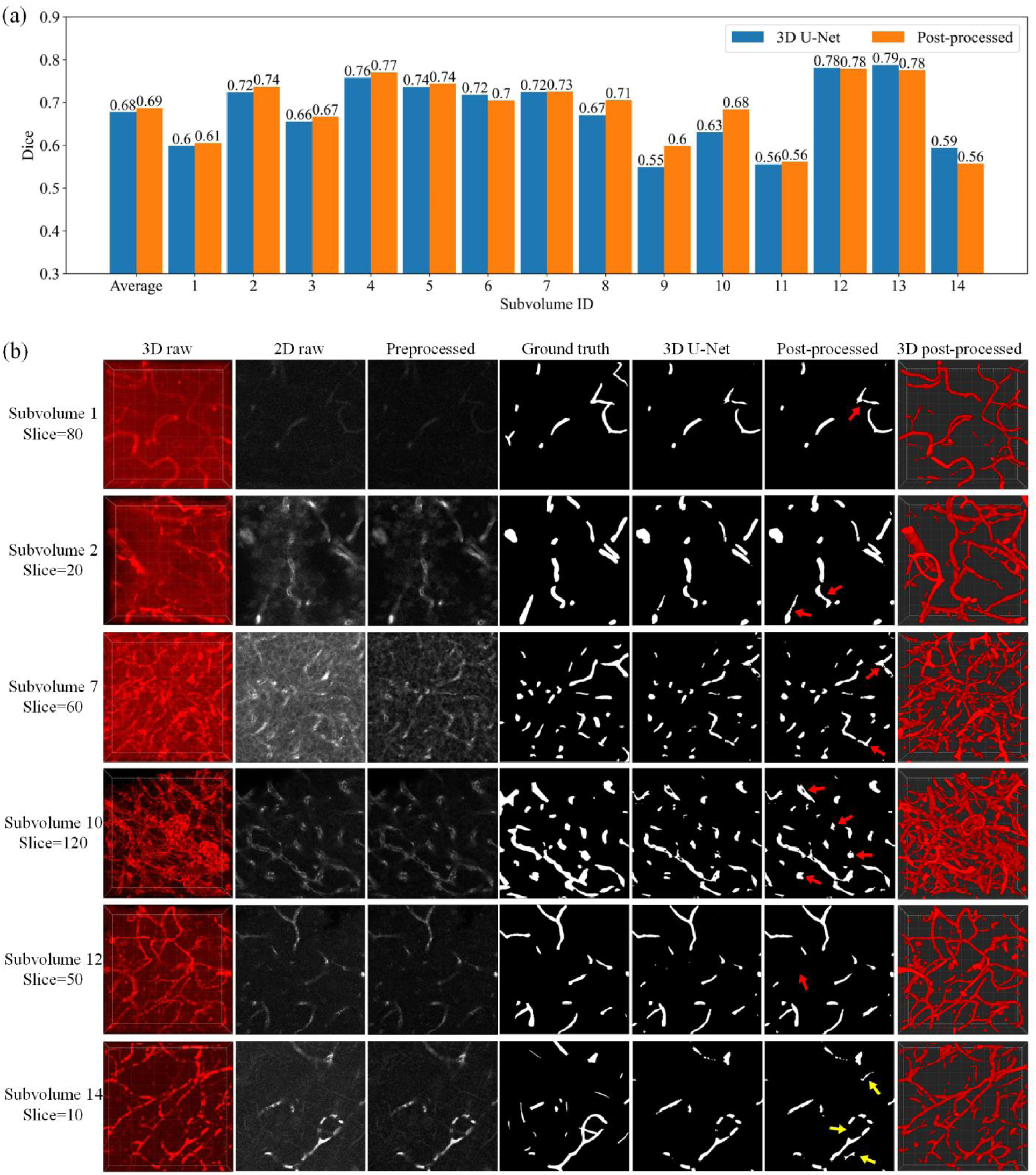
Vessel extraction results with our deep learning-based algorithms. (a) The Dice values of all the 14 subvolumes before and after the post-processing step. Subvolume 1 to 8 are from the low-grade gliomas and subvolume 9 to 14 are from the high-grade; (b) Visualization of representative extracted vessels. The results of 6 representative subvolumes in three steps, including preprocessing, 3D U-Net segmentation, and post-processing, are presented. In the post-processing step, the red arrows point out the main correct improvements and the yellow show the incorrect cases.

In most cases, the Dice value was not high but the 3D visualization performed well, due to the fact that the vessels travel in 3D space and the singe slice can not capture the complete vessels, which makes the manual annotation of slices ambiguous. Subvolume 14 is a representative data with a low Dice value and the reason for the bad results is that the vessels are quite discontinuous while those in label images are continuous. In addition, the Dice value decreases after the post-processing step (Figure 5 a). The reason is that the small vessel segments are prone to be treated as noise and removed incorrectly, and some neighbouring vessels are incorrectly connected (See yellow arrows in Figure 5 b). Overall, our 3D U-Net achieved reliable segmentation results and subsequent post-processing methods finetuned the results. Finally, we obtained a segmentation mask containing intact tumor vessels and a clean background.

In addition, we applied the segmentation methods to the whole volumes and compared the results with those of Imaris (Figure 6). The overlap between the raw image and extracted vessels was shown to visualize the vessel extraction performance. Obviously, our deep learning methods performed much better than Imaris. In the 3D raw images, we can see the glioma vessels have various morphology and scale with varying degrees of breakage and blurring, and the images have varied brightness, contrast, and background interference. Compared with Imaris, deep learning based methods can cope with the complex glioma vessels with a better quality. Specifically, it can detect both tiny and large vessels with varied contrast and brightness (e.g. Figure 6 d-f). The intact vessel segments can be extracted well although the segments are broken and discontinuous (e.g. Figure 6 c, e). Furthermore, it enables better differentiation of autofluorescent spots and discrete tiny tumor vessel segments than the Imaris (e.g. Figure 6 b). For the images with fuzzy vessels and strong background, which is even difficult for human experts, our model can also segment the vessels based on the experience learnt by the neural network (Figure 6 c). Hence, compared with Imaris, our deep learning method is more accurate and fully automated in the high-throughput and bias-free vessel extraction.

**Fig. 6.**
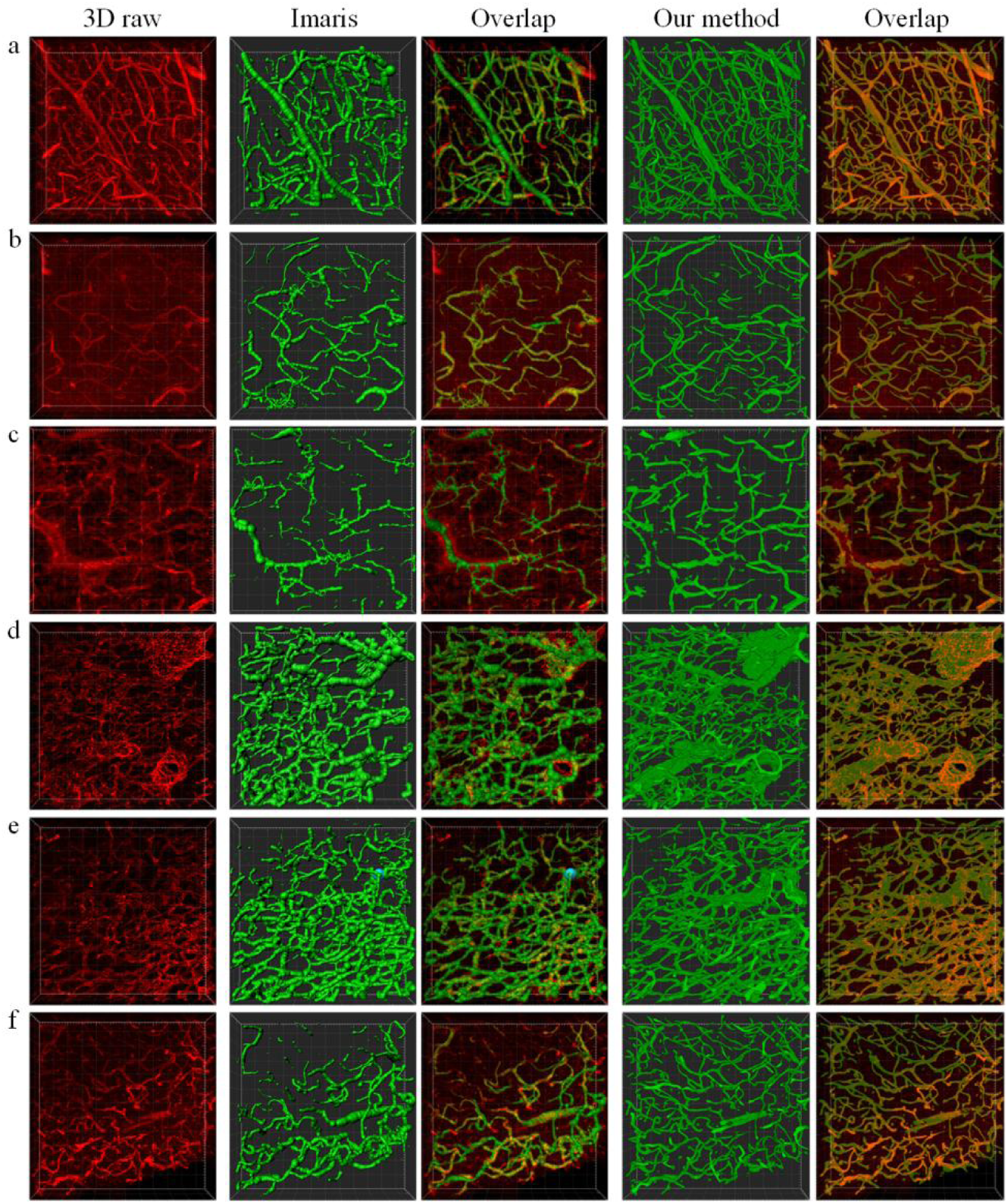
Comparison of 3D vessel extraction between the Imaris and our deep learning based methods. Representative volumes were shown. Volume a-c are from the low-grade gliomas and volume d-f are from the high-grade. Vascular immunofluorescence volumes are shown in red. Imaris results and 3D U-Net segmentation masks are shown in green. The overlap images represent the extraction performance.

### 3.4 Quantitative analysis of vascular morphological features reveals vascular heterogeneity in gliomas

Based on the segmentation masks, we extracted and calculated the vascular morphological features and performed quantitative analysis. For three groups, namely normal, low-grade, and high-grade gliomas, we selected the volumes with a better quality of vascular visualization for reliable quantitative analysis. As a result, there were 10, 17 and 12 volumes in there groups. They are from the samples of 2 patients with normal brain, 6 patients with low-grade glioma, and 2 patients with high-grade glioma, respectively.

Figure 7 shows the quantitative analysis results. Significant heterogeneity in the structure of blood vessels between different grades of gliomas was observed. Firstly, the mean diameter of blood vessels in low-grade gliomas is 6.34 μm while in high-grade gliomas is 7.59 μm, demonstrating that the vessel diameters in high-grade gliomas were significantly larger than those in low-grade gliomas (p = 0.0156). Secondly, the average number of vascular branching points per unit volume in low-grade gliomas is 30572 and in high-grade gliomas is 57876, suggesting that high-grade gliomas tend to generate more branches (p = 0.0295). Thirdly, the average volume ratio, the surface area ratio of vessels, and the average vessel length in per unit volume of high-grade gliomas are all larger than those in low-grade gliomas (p = 0.0209, 0.0457, 0.0161). These observations reflect the larger vascular density in high-grade gliomas. However, we also noted that there are more branches and longer average length of blood vessels in normal brain tissue than in low-grade gliomas, which may be attributed to the influence of discontinuous staining of broken vessels.

**Fig. 7.**
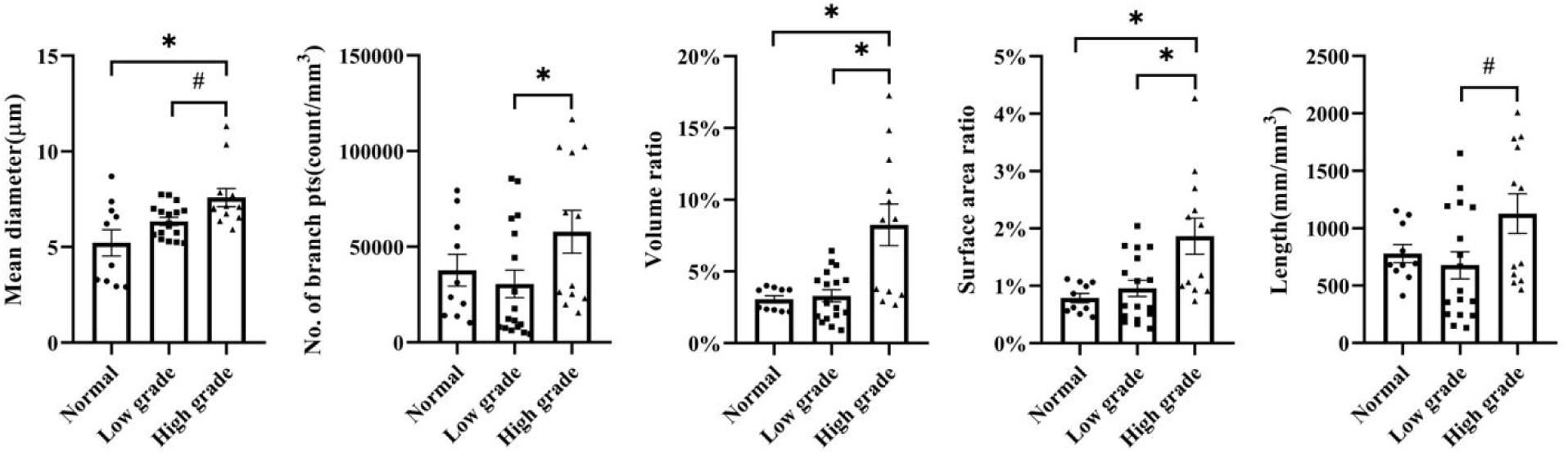
Quantitative analysis of vascular morphological features in normal brain tissue, low-grade, and high-grade gliomas. Mean diameter: mean diameter along the full length. No. of branch pts: number of branching points per unit volume. Volume ratio: ratio of the total voxel number of vascular volume to the total voxel number of the whole volume. Surface area ratio: ratio of the total voxel number of vascular surface to the total voxel number of the whole volume. Length: total vessel length per unit volume. *p < 0.05, **p < 0.01, ***p < 0.001, ****p < 0.0001, muti-group comparisons. #p < 0.05, ##p < 0.01, ###p < 0.001, ####p < 0.0001, two-group comparisons.

## 4 Discussion

High-grade gliomas, like most tumors, have a high degree of vascular heterogeneity in advanced lesions [25]. However, 2D based analysis methods suffer from structural overlap, particularly in highly heterogeneous gliomas, where paraffin sections separated by tens of micrtons can reflect different vascular characteristics, and even the structure of vessels in different sites of the same surface can vary considerably. In contrast to the routine neuropathological examination performed on thin tissue sections, our pathoanatomical study of thick tissue sections achieved 3D visualization of vessels in large scale human glioma tissue blocks (5000μm×5000μm×500μm) at cellular resolution, which complemented conventional histology in terms of 3D spatial imaging and pathological heterogeneity assessment of intact tissue. Based on the 3D images, spatial morphological heterogeneity in different grades of gliomas can be visually observed.

With the help of deep learning, data analysis based on tissue clearing technology will evolve towards high accuracy, high throughput, bias-free and automation. Deep learning method is the best solution for extracting complex tumor vessels, especially glioma vasculature with high heterogeneity. In addition, its high performance can compensate for the shortcomings of data acquisition techniques. To the best of our knowledge, we are the first to use an end-to-end deep neural network for segmenting tumor vessels in cleared tissue images. The architecture of 3D U-Net, a variety of data augmentation, larger size of input images, and a higher number of iterations during the training are key to obtaining high extraction performance. For our glioma vessels with higher degree of fragmentation, a higher iteration number of closing operation could help obtain their more intact masks. We considered that tumor vessels with intact shapes can be easily quantified using quantitative criteria for normal vessels, which allows for a more rational comparison of quantitative metrics. Due to the limited amount of reliable data and the large variations in different volumes of the same sample, we performed the statistical analysis at the volume level. By comparing low-grade (WHO grade II) with high-grade (WHO grade IV) glioma vessels, we found that high-grade glioma vessels tend to generate more vascular branches, exhibit larger mean vessel diameters, and having larger volume, surface area and total length than low-grade glioma vessels. The reasonableness of this findings is supported by the experimental results of Lagerweij et al. based on animal models[21].

Since the diameter of blood vessels is closely related to the properties of hemodynamics, a larger vessel diameter implies less resistance to blood flow and a higher number of branches implies having a larger amount of neovascularization[44, 45]. Accordingly, we might speculate that high-grade gliomas have a higher demand for blood than low-grade gliomas. This may be related to the fact that high-grade gliomas need more nutrients and oxygen to support their strong aggressiveness and high resistance. Since leakage of the vascular wall can cause deregulated permselectivity and perfusion, high interstitial fluid pressure, extensive hypoxia, necrosis, and edema[18], and hypoxia can alter gene expression in tumor cells, thereby increasing cell survival and resistance to apoptosis induction [54], it can be speculated that high-grade gliomas with poorer vascular wall integrity benefit more from glioma cell resistance based on vascular leakage mechanisms. This firmly supports the hypothesis that high-grade gliomas are more dependent on blood vessels compared to low-grade gliomas. Thus, theoretically, high-grade gliomas may show better responsiveness to vascular-targeted therapy. Based on this mechanistic conjecture, a series of studies targeting VEGF for anti-angiogenic treatment of GBM are underway. However, there is insufficient evidence to support that this strategy significantly improves survival in patients with glioblastoma[2, 5, 12]. The limited availability of anti-VEGF strategies may be related to the existence of other mechanisms for brain tumor angiogenesis, including endothelial differentiation of glioblastoma stem-like cells[16, 43, 52], and vascular co-option[16, 40]. In recent years, exploring macrophage-associated immunosuppression to regulate glioblastoma angiogenesis[8], glial cell signaling in vascular co-selection[14], and the influence of other tumor microenvironment components on glioma angiogenesis has opened new avenues for research into vascular-targeting therapies. Leveraging 3D pathology enabled by tissue clearing technology allows researchers to investigate the impact and mechanisms of the tumor microenvironment on blood vessels within expansive tissue blocks, facilitating the identification of novel targets for vascular-targeted glioma treatments[55].

It is worth noting that our study is preliminary and has some key limitations. While we have not found evidence of uneven labeling and changed diffusivity in our samples, we cannot rule these out as possibilities. Although our results of quantitative analysis were consistent with vision observation and previous studies, more careful design, more rigorous execution, and more samples are needed for validation further. After compensating for these limitations, we are able to do further research on the glioma vasculature. In the future, we would improve the methods of tissue clearing for more data with higher quality. Meanwhile, vessels extraction methods would be improved for higher accuracy. Based on the extracted vessels, we would explore more morphological features characterizing vascular heterogeneity. Based on enough glioma vessel data, we will perform more reliable quantitative analysis in the patient level. Finally, each step will be optimized and a platform for glioma vasculature analysis based on tissue clearing and deep learning will be built. By continuously enriching sample size and correlating it with other microenvironmental components and clinical information, our protocol and platform will help to assist in the graded diagnosis of glioma and the establishment of new criteria for prognostic assessment[30, 50, 58], and achieve precise treatment of subpopulations of glioma cells with vascular heterogeneity.

## 5 Conclusions

We demonstrate the use of tissue clearing and deep learning technology to study in 3D the architecture of glioma vessels at cellular resolution and to assess the spatial heterogeneity of glioma vessels, which can complement conventional histology in 3D spatial imaging and in assessing the pathological heterogeneity of intact tissues. Our study provided an experimental protocol for 3D reconstruction, qualitative and quantitative analysis of human glioma vascular heterogeneity, which may bring value to basic and clinical research on glioma vasculature and microenvironment.

## Abbreviations

2D: two-dimensional;
3D: three-dimensional;
BBB: blood-brain barrier;
DAPI: 4 ’ 6-diamidino-2-phenylindole;
FFPE: formalin-fixed, paraffin-embedded;
GBM: glioblastoma;
NBF: neutral buffered formalin;
PBS: phosphate-buffered saline;
SBC: Sodium borate clearing;
TME: tumor microenvironment;
ECM: extracellular matrix;
EV: extracellular vesicles;
BBB: blood-brain barrier;
TDE: 2,2 ’ -thiodiethanol;
PBST: phosphate-buffered saline with Triton X-100.

## Declarations

### Ethical approval and consent to participate

The tissue used in this study was provided by the Clinical Biobank Center at Zhujiang Hosipital of Southern Medical University. The research protocol has been approved by the Medical Ethics Committee of Zhujiang Hospital of Southern Medical University (Approval Number: 2018-SJWK-004 and 2020-YBK-001-02) and informed consent was obtained from all patients.

### Consent for publication

Not applicable.

### Availability of data and materials

The code of deep learning is available at https://github.com/PRBioimages/vesselsegmentation, and the data used in manuscript is available upon reasonable request to the corresponding authors.

### Authors’ contributions

XW: Conceptualization, Formal Analysis, Investigation, Methodology,Writing - Original Draft preparation, Writing - Review & Editing. XY: Formal Analysis, Algorithm development, Writing - Review & Editing. DH: Formal Analysis, Investigation, Methodology, Writing - Review & Editing. YL: Investigation, Methodology, Writing - Review & Editing. QH: Investigation, Methodology, Writing - Review & Editing. TL: Investigation, Methodology, Writing - Review & Editing. ZY: Investigation, Writing - Review & Editing. CY: Investigation, Writing - Review & Editing. MZ: Investigation, Writing - Review & Editing. HL: Technical expertise, Writing - Review & Editing. YX: Conceptualization, Supervision, Funding acquisition, Project administration, Writing - review & editing. HS: Conceptualization, Supervision, Funding acquisition, Project administration, Writing - review & editing.

## Acknowledgements

Not applicable.

